# Motor Cortical Neuronal Hyperexcitability Associated with α-Synuclein Aggregation

**DOI:** 10.1101/2024.07.24.604995

**Authors:** Liqiang Chen, Hiba Douja Chehade, Hong-Yuan Chu

**Author notes:** Correspondence: Dr. Hong-Yuan Chu.

## Abstract

Dysfunction of the cerebral cortex is thought to underlie motor and cognitive impairments in Parkinson disease (PD). While cortical function is known to be suppressed by abnormal basal ganglia output following dopaminergic degeneration, it remains to be determined how the deposition of Lewy pathology disrupts cortical circuit integrity and function. Moreover, it is also unknown whether cortical Lewy pathology and midbrain dopaminergic degeneration interact to disrupt cortical function in late-stage. To begin to address these questions, we injected α-synuclein (αSyn) preformed fibrils (PFFs) into the dorsolateral striatum of mice to seed αSyn pathology in the cortical cortex and induce degeneration of midbrain dopaminergic neurons. Using this model system, we reported that αSyn aggregates accumulate in the motor cortex in a layer- and cell-subtype-specific pattern. Particularly, intratelencephalic neurons (ITNs) showed earlier accumulation and greater extent of αSyn aggregates relative to corticospinal neurons (CSNs). Moreover, we demonstrated that the intrinsic excitability and inputs resistance of αSyn aggregates-bearing ITNs in the secondary motor cortex (M2) are increased, along with a noticeable shrinkage of cell bodies and loss of dendritic spines. Last, neither the intrinsic excitability of CSNs nor their thalamocortical input was altered by a partial striatal dopamine depletion associated with αSyn pathology. Our results documented motor cortical neuronal hyperexcitability associated with αSyn aggregation and provided a novel mechanistic understanding of cortical circuit dysfunction in PD.

## Introduction

Degeneration of dopaminergic neurons in the substantia nigra pars compacta (SNc) has been associated with accumulation of cytoplasmic Lewy-like pathology in most PD cases (Spillantini et al., 1997). Decreased levels of dopamine (DA) in the basal ganglia significantly alter the connectivity and computation of the basal ganglia-thalamocortical circuits in parkinsonism. Traditional circuit model of PD pathophysiology suggests that the aberrant inhibitory output of the basal ganglia disrupts the thalamic excitation to the motor cortex, which decreases cortical motor output and underlies the hypokinetic symptoms in PD (Albin et al., 1989; DeLong, 1990; Galvan and Wichmann, 2008; McGregor and Nelson, 2019). This model assumes that cortical microcircuits remain intact in parkinsonian state and that cortical hypofunction is an instant effect of the exaggerated suppression of thalamus by the basal ganglia. However, converging evidence from recent studies suggest that the motor cortex shows intrinsically disrupted connection and function in animal models of parkinsonism that can play a major role in cortical pathophysiology of PD (Chu, 2024). These local cortical changes include altered dendritic spine dynamics and density of cortical pyramidal neurons and reduced thalamic axonal markers in the primary motor cortex (M1) of both parkinsonian monkeys and rodents (Guo et al., 2015; Villalba et al., 2019; Chen et al., 2023). Such anatomical changes also associate with altered thalamocortical connection strength and dampened intrinsic excitability of pyramidal tract neurons (PTNs) in M1 (Chen et al., 2021; Chen et al., 2023; Swanson et al., 2023). Consistently, *in vivo* studies also reported a disrupted timing and magnitude of M1 PTNs activation in mediating motor activities in parkinsonian animals (Pasquereau and Turner, 2011; Pasquereau et al., 2016; Aeed et al., 2021). Thus, the motor cortex is a site of intrinsic dysfunction, instead of being an information gateway that translates basal ganglia abnormalities into motor deficits in PD.

Most prior research on cortical dysfunction has been conducted using dopamine-depleted neurotoxin models of parkinsonism that usually do not develop Lewy-like pathology. Post-mortem studies of human PD reported moderate levels of Lewy pathology in cerebral cortical motor regions at the Braak stages 4-6, indicating a potential role of cortical pathology in motor and cognitive impairments in PD (Hurtig et al., 2000; Braak et al., 2003; Dickson et al., 2010; Fu et al., 2022). However, how the development of Lewy pathology disrupts motor cortical circuit integrity and function remains largely unexplored. In the present the study, we studied the functional impact of α-synuclein (αSyn) aggregates on motor cortical neurons and their synaptic inputs using αSyn preformed fibrils (PFFs)-seeding model of synucleinopathy. We demonstrated that αSyn aggregates triggered hyperexcitability and shrinkage of intratelencephalic neurons (ITNs) in the secondary motor cortex (M2). Moreover, we found that mild loss of striatal dopamine associated with αSyn aggregation is not sufficient to induce changes in the intrinsic and synaptic properties of corticospinal neurons (CSNs) in M2. These results demonstrate that the intrinsic excitability of cortical pyramidal neurons are enhanced by the accumulation of intracellular αSyn aggregates and that severe striatal dopamine loss is required to induce adaptive circuit changes in the motor cortex in parkinsonism.

## Materials and methods

### Animals

Three-to-four months old wild type male C57BL/6J mice (JAX stock#:000664, RRID: IMSR_JAX:000664) were used in this study and were provided by the Van Andel Research Institute vivarium. Mice were housed up to four animals per cage under a 12h/12h light/dark cycle with free access to food and water in accordance with NIH guidelines for animal care and use. Animal studies were reviewed and approved by the Institutional Animal Care and Use Committee (IACUC) at Van Andel Institute (animal use protocol #: 22-02-007).

### Preparation and validation of **α**Syn preformed fibrils

Escherichia coli BL21 codon plus RIPL cells (RRID: CVCL_M639) were used to produce and purify mouse αSyn protein, which was then dialyzed using a buffer containing 10 mM Tris and 50 mM NaCl (pH 7.5). Endotoxins were removed using a high-capacity endotoxin removal kit (PI88276) and then were assessed using an endotoxin quantification kit (A39552). The protein concentration was estimated using absorbance at 280 nm with an extinction coefficient of 7450 M^−1^ cm^−1^. Purified mouse αSyn monomer protein was used to generate mouse αSyn preformed fibrils. Specifically, monomeric αSyn protein was diluted to 5 mg/mL in the buffer (150 mM KCl and 50 mM Tris-HCl), incubated at 37°C with shaking for 7 days, and centrifuged for 10 min at 13,200 rpm (Volpicelli-Daley et al., 2014). The protein pellet was re-suspended in half of the initial volume of the solution.

Fibril solution (5 μl) was incubated with 5 μl of 8 M guanidinium chloride at room temperature for one hour and the concentration of PFFs was measured using absorbance at 280 nm. PFFs were diluted at 5 mg/mL and 22-25 μl aliquots were stored at −80°C until use. On the day of injection, an aliquot of PFFs (22-25 μl at 5 mg/mL) was thawed at room temperature and sonicated using Qsonica 700W cup horn sonicator at 30% amplitude using 3 seconds on/2 seconds off cycle for 15 min at 15°C. The size of sonicated PFF (30-70 nm segments) was estimated and confirmed using the dynamic light scattering (DynaPro NanoStar from Wyatt technology). Details of generation and validation of αSyn PFFs can be found here: dx.doi.org/10.17504/protocols.io.bhhrj356.

### Stereotaxic brain surgery

Isoflurane (2-3%) was used to induce anesthesia. Mice were placed into in a stereotaxic frame (Stoelting, Model: 51730M) with head fixed using ear bars. A feedback controller was used to maintain and monitor the body temperature. To induce αSyn pathology in the motor cortex, sonicated PFFs (2 µl at 5 µg/µl) were delivered into the dorsolateral striatum [from bregma in mm, anteroposterior (A-P) = +0.2, mediolateral (M-L) = −2.3, dorsoventral (D-V) = −2.9] through a glass pipette attached to a Nanoliter injector (NANOLITER-2020, World Precision Instrument, FL, USA) with a rate of 0.4 µl per minute. Injection glass pipettes were made by a vertical pipette puller (Model 720, David Kopf Instruments, CA, USA). Mice receiving αSyn monomers or phosphate-buffered saline (PBS) were used as controls. Retrobeads (1:10 dilution, Lumafluor) were mixed with PFFs or monomers or PBS, as appropriate, and co-injected into the dorsolateral striatum or the spinal cord (C7-8) to retrogradely label ITNs or CSNs, respectively. To study thalamocortical transmission using optogenetics, AAV9-hSyn-ChR2(H134R)-eYFP (Addgene#127090, RRID:Addgene_127090; volume = 300 nl, titer = 3.6 × 10^12^ GC/ml) were stereotaxically injected into the motor thalamus centered at the ventromedial subregions (from bregma in mm, A-P = −1.3, M-L = +0.75, D-V = −4.3). Stereotaxic brain surgery procedure details can be found on Protocols.io (dx.doi.org/10.17504/protocols.io.rm7vzye28lx1/v1).

### Spinal cord surgery

Mouse spinal cord surgery was performed as described previously (Chaterji et al., 2021). Mouse was placed into a stereotaxic frame (Stoelting) and anesthetized using 2-3% isoflurane. Body temperature was maintained and monitored using a heating pad connected to a feedback controller. An incision (1 cm) was made over the spinal cord and vertebrae were exposed using retractors. Once the C7 and C8 segments were located, Retrobeads and/or PFFs (5 µg/µl, 1 µL/segment) were injected into the spinal cord (500 µm away from the midline, 700 µm in depth) via a glass pipette mounted on a Nanoliter injector (NANOLITER-2020, WPI, Florida, USA) at a rate of 0.2 µl per minute. Details of mouse spinal cord injections can be found on protocols.io: dx.doi.org/10.17504/protocols.io.81wgbz5e3gpk/v1

### Preparation of brain slices for electrophysiology

Mouse was anesthetized using avertin (300 mg/kg) and perfused transcardially using ice-cold sucrose-based cutting solution containing 230 mM sucrose, 26 mM NaHCO_3_,10 mM glucose, 10 mM MgSO_4_, 2.5 mM KCl, 1.25 mM, NaH_2_PO_4_, 0.5 mM CaCl_2_, 1 mM sodium pyruvate, and 0.005 mM L-glutathione. Mouse brain was carefully dissected and coronal brain sections (250 µm in thickness) containing the motor cortex were prepared using a VT1200S vibratome (Leica Microsystems Inc., Deer Park, IL; RRID:SCR_018453). Brain sections were then kept in normal artificial cerebrospinal fluid (aCSF) containing 126 mM NaCl, 26 mM NaHCO_3_,10 mM glucose, 2.5 mM KCl, 2 mM CaCl_2_, 2 mM MgSO_4_, 1.25 mM NaH_2_PO_4_,1 mM sodium pyruvate, and 0.005 mM L-glutathione. Slices were incubated in aCSF at 35°C for 30 min, and then kept at room temperature until recording. Details of brain section preparation can be found on Protocols.io (dx.doi.org/10.17504/protocols.io.36wgqj2eovk5/v1).

### *Ex vivo* electrophysiology recording

Brain sections were transferred to a recording chamber perfused with synthetic interstitial fluid (SIF) containing 126 mM NaCl, 26 mM NaHCO_3_,10 mM glucose, 3 mM KCl, 1.6 mM CaCl_2_,1.5 mM MgSO_4_, and 1.25 mM NaH_2_PO_4_ at a rate of 3.5 mL/min. SIF solution was equilibrated with 95% O_2_ and 5% CO_2_ and maintained at 33-34°C via a feedback controlled in-line heater (TC-324C, Warner Instruments). Neurons were visualized by a 60x water immersion objective lens (Olympus, Japan) using SliceScope Pro 6000 system integrated with a charge-coupled device camera (SciCam Pro, Scientifica, UK). Whole-cell patch-clamp recording was performed using a MultiClamp 700B amplifier (Molecular Devices, San Jose, CA; RRID:SCR_018455) and Digidata 1550B controlled by pClamp 11 software (Molecular Devices, San Jose, CA; RRID:SCR_011323). Glass pipettes (BF150-86-10, Sutter Instruments,) were prepared using a micropipette puller (P1000, Sutter Instruments, Novato, CA, USA; RRID:SCR_021042) and filled with one of the following internal solutions: (1) K-gluconate-based internal solution (140 mM K-gluconate, 3.8 mM NaCl, 1 mM MgCl_2_, 10 mM HEPES, 0.1 mM Na_4_-EGTA, 2 mM ATP-Mg, and 0.1 mM GTP-Na pH = 7.3, mOsm = 290); or (2) Celsium-methanesulfonate-based internal solution (120 mM CH_3_O_3_SCs, 2.8 mM NaCl, 10 mM HEPES, 0.4 mM Na_4_-EGTA, 5 mM QX314-HBr, 5 mM phosphocreatine, 0.1 mM spermine, 4 mM ATP-Mg, and 0.4 mM GTP-Na, pH = 7.3, mOsm = 290). Biocytin (0.2%) was included into the K-gluconate internal solution to label the recorded neurons for morphology studies. Ionotropic glutamatergic and GABAergic synaptic transmission was blocked by a cocktail of DNQX (20 μM), D-APV (50 μM), and SR-95531 (10 μM) in the recording solution for intrinsic excitability recording. TTX (1 µM) and 4-AP (100 µM) were routinely included in the recording solution to isolate monosynaptic thalamocortical transmission. Details of *ex vivo* electrophysiological recording can be found on: Protocols.io (dx.doi.org/10.17504/protocols.io.eq2ly7m2rlx9/v1).

### Immunofluorescent staining

#### Immunohistology of αsyn pathology and biocytin

After electrophysiology recording, the brain sections (250 µm) containing biocytin-filled neurons were placed into 4% paraformaldehyde (PFA) solution overnight at 4°C. Brain sections were then rinsed for three times using phosphate-buffered saline (PBS), and incubated with 2% normal donkey serum in 0.5% Triton-X-100 PBS solution for 1 hour, followed by incubation with primary antibody against pS129 αSyn (Rabbit, 1:10,000, Bioscience, #1536-1, RRID: AB_562180) overnight at room temperature. The brain sections were rinsed for three times with PBS and incubated with the secondary antibody AlexaFluor 488 donkey anti-rabbit IgG (1:500, cat#: 711-545-152, Jackson ImmunoResearch Labs, RRID: AB_2313584) and Cy5-conjugated streptavidin (1:1000, cat#: SA1011, Thermo Fisher Scientific) for two hours at room temperature. After 3x rinse with PBS, brain sections were mounted on slides using mounting medium (H-1000, Vector Laboratories) and cover slipped.

#### Immunohistology of tyrosine hydroxylase (TH)

Remaining brain tissue from slice preparation for electrophysiology was immersed in 4% PFA solution overnight at 4°C, and re-sectioned (70 µm) using a VT1000s vibratome (Leica Biosystems, Deer Park, IL; RRID:SCR_016495). Slices containing the striatum and the substantia nigra pars compacta were collected for tyrosine hydroxylase (TH) or/and phospho-Ser129 (pS129) αSyn immunohistochemistry using the following primary and secondary antibodies: primary antibodies [mouse anti-TH (1:2000, cat#: MAB318, MilliporeSigma; RRID: AB_2201528) and rabbit anti-pS129 α-Syn (1:10,000, Bioscience, cat#1536-1, RRID: AB_562180)] and secondary antibodies [Alexa Fluor 488 donkey anti-mouse IgG (1:500, cat#715-545-150; Jackson ImmunoResearch Labs, RRID: AB_2340846) and Alexa Fluor 647 donkey anti-rabbit IgG (1:500, cat#711-605-152, Jackson ImmunoResearch Labs, RRID: AB_2492288)]. Details of immunofluorescent staining can be found on Protocols.io (https://www.protocols.io/view/immunofluorescent-staining-3byl4bq9ovo5/v1).

#### Quantification of αSyn pathology in the motor cortex

The coronal sections of the motor cortex with the following A-P coordinates from the bregma (in mm): +1.94, +1.70, +1.54, and +1.18, were used for quantification of αSyn pathology. The brain sections were manually aligned to the mouse brain atlas of Franklin and Paxinos (5^th^ Edition, 2019; ISBN: 9780128161579) to recognize M1 and M2. The cortical layer boundaries were determined based on a previous report (Oswald et al., 2013): layer 1 (0 - 0.1 mm from the pia), layer 2/3 (0.1 - 0.4 mm), layer 5A (0.4 - 0.55 mm), layer 5B (0.55 - 0.8 mm), and layer 6 (0.8 - 1.0 mm). ImageJ (NIH, https://imagej.net/, RRID: SCR_003070) was used to quantify the proportion of areas occupied by pS129 αSyn pathology, as reported in our recent work (Chen et al., 2022; Zhou et al., 2024).

### Confocal imaging

Immunofluorescent images were collected using a Nikon A1R Confocal Laser Scanning Microscope (RRID:SCR_020317). pS129 αSyn pathology in cortical area were imaged under 20x objective lens and quantified in ImageJ (NIH, https://imagej.net/, RRID: SCR_003070). Biocytin-filled cortical neurons were imaged under 20x lens (NA=0.75, x/y, 1024/1024 pixels, z-step=1 µm). Spines of biocytin-filled cortical neurons were imaged under a 100x objective lens (NA=1.45, x/y, 1024/1024 pixels, z-step = 0.5 µm). Spine density was assessed from 2-3 segments of basal dendrites (20-30 µm in length) at a distance between 50 and 100 µm measured from the soma, which were reconstructed manually using the filament tracer function of Imaris software (Version 10.1.1, Oxford, UK, http://imaris.oxinst.com, RRID: SCR_007370). Details of confocal imaging can be found on Protocols.io: https://www.protocols.io/view/confocal-imaging-and-digital-image-analysis-3byl4jmxzlo5/v1.

### Data analysis and statistics

Electrophysiology data were analyzed using Clampfit software (Version 11.1, Molecular Devices, San Jose, USA, RRID: SCR_011323). The amplitude of EPSCs in response to blue light stimulation was quantified to measure synaptic connection strength. Confocal images were analyzed using Imaris (Version 10.1.1, Oxford, UK, http://imaris.oxinst.com, RRID: SCR_007370) or ImageJ (NIH, https://imagej.net/, RRID: SCR_003070) for spine density quantification or Sholl analysis, respectively. GraphPad Prism (Version 10, GraphPad Software, http://www.graphpad.com, RRID: SCR_002798) was used for statistics analysis. Non-parametric, distribution-independent Mann-Whiney U (MWU) test was used to compare the median of two groups and Kruskal-Wallis test was used to compare the median three or more groups. Two-way analysis of variance (ANOVA) was used to compare the main effects of group difference in the amplitude of EPSCs or the frequency of action potentials across a range of stimulation intensities, followed by Dunn’s multiple comparisons test. *P* < 0.05 was considered as statistically significant.

## Results

### Layer- and cell-type-specific pattern of αSyn pathology distribution in the motor cortex of intrastriatal PFFs-injected model

In the intrastriatal PFFs seeding model, αSyn pathology reaches and stays at peak levels in rodent brain at 3 months post-injection (mpi) (Luk et al., 2012; Burtscher et al., 2019; Henderson et al., 2019; Stoyka et al., 2019). Thus, all studies in the following sections were conducted at 3 mpi when cortical pathology was robust, unless stated otherwise.

We detected robust phosphorylated αSyn at serine129 (pS129)-immunoreactive (ir) aggregates, an indicative of pathologic αSyn, in the primary and secondary motor cortices of both hemispheres (M1 and M2, respectively) at 3 mpi (**Figure 1A**). No pS129-ir αSyn aggregates were detectable in the cerebral cortices of either PBS- or αSyn monomer injected mice (**Supplementary Figure 1A**), as reported previously (Luk et al., 2012; Stoyka et al., 2019; Gcwensa et al., 2024). Moreover, the level of αSyn pathology was higher in the M2 than the M1 in both hemispheres (**Figure 1B**). In M2, pS129-ir αSyn aggregates were not evenly distributed across cortical layers; instead, they showed highest levels in the layer 5A, followed by the layer 6, layer 5B, layers 2/3 and 1 (**Figure 1C**, see also (Goralski et al., 2024)).

**Figure 1.**
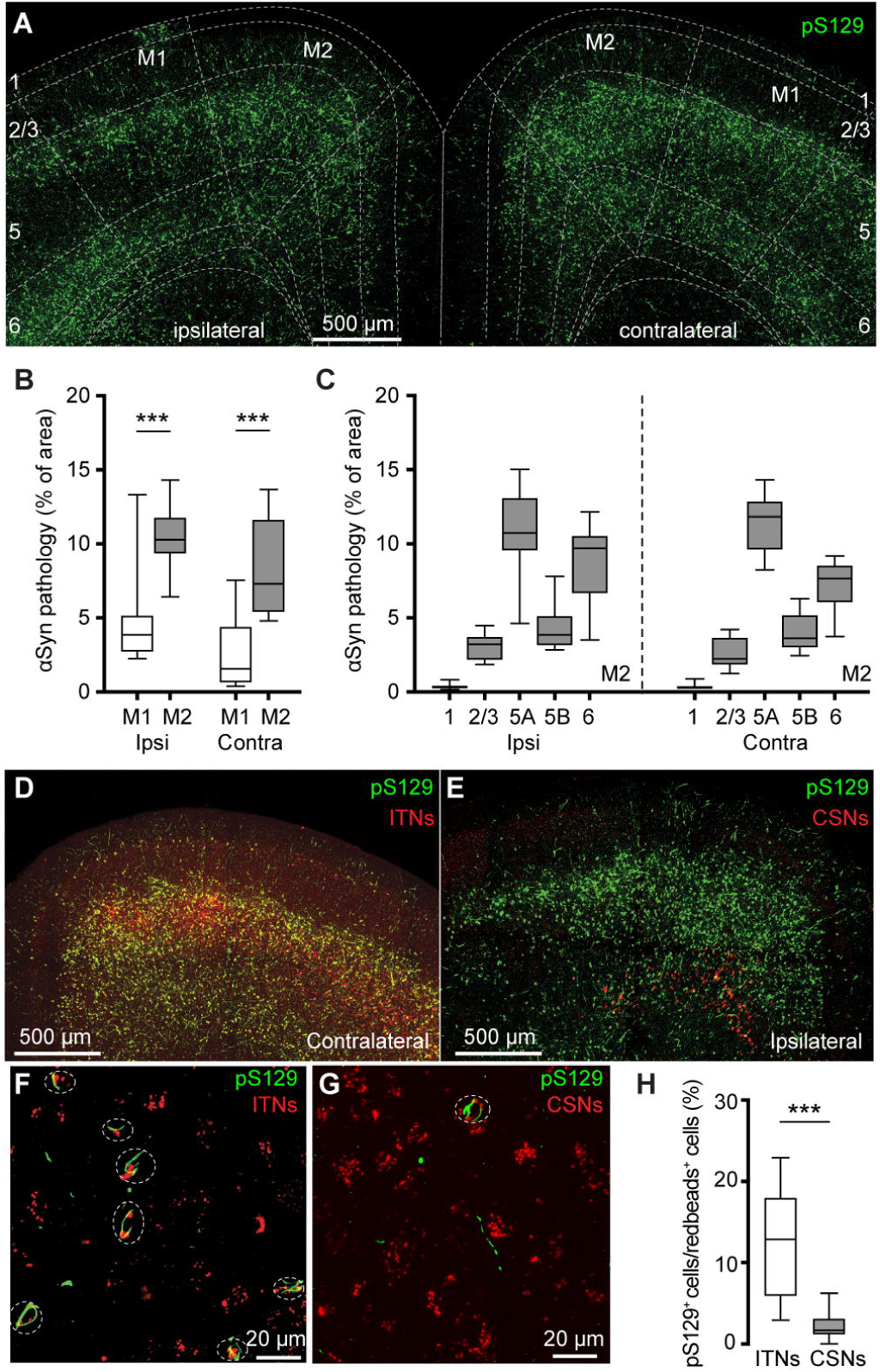
PFFs injection into the dorsal striatum seeds αSyn pathology in motor cortex. **(A)** Representative confocal images showing the pS129^+^ αSyn pathology in the motor cortex at 3 months-post-injection (mpi). **(B)** The proportion of the motor cortical area covered by the pS129^+^ αSyn aggregation. (Ipsilateral, M1 = 3.9 [2.7, 5.2]%, M2 = 10.3 [9.3, 11.8]%, n = 12 slices/4 mice for each group; *p* = 0.0005, Mann–Whiney U (MWU) test. Contralateral, M1 = 1.6 [0.6, 4.4]%; M2 = 7.3 [5.4, 11.7]%, n = 12 slices/4 mice for each group, *p* = 0.0001, MWU). **(C)** The proportion of the different cortical layers covered by pS129^+^ αSyn aggregation. **(D-G)** Representative images showing the co-localization of Redbeads labeling with pS129^+^ αSyn pathology in motor cortex at low (D-E) and high (F-G) magnifications. (**H**) Summarized results showing that a higher percentage of ITNs than CSNs in M2 were pS129^+^ αSyn positive (ITNs = 12.9 [5.9, 18.0]%, n = 11 slices/4 mice; CSNs = 1.7 [1.1, 3.2]%, n = 12 slices/4 mice; *p* < 0.0001, MWU).

In PFFs-based models, development of αSyn pathology in the cortex involves the uptake of PFFs at the corticostriatal axonal terminals (Volpicelli-Daley et al., 2011; Luk et al., 2012). Thus, the layer-specific accumulation of αSyn aggregates in M1/M2 is consistent with the anatomical and physiological studies showing that corticostriatal projection neurons are mainly found in the layers 5 in rodent brains (Wilson, 1987; Levesque et al., 1996; Shepherd, 2013).

Since the M2 showed robust pathology, we further interrogated the effects of αSyn aggregation on its microcircuits. Intratelencephalic neurons and corticospinal neurons (a subset of PTNs) are mainly located in the layers 5A and 5B of M2, respectively. Considering the striking difference in the amount of pathology between layer 5 subregions, we tested the hypothesis that ITNs develop heavier αSyn aggregation than CSNs. To estimate the proportion of aggregates-bearing ITNs and CSNs, we unilaterally injected (1) Retrobeads into the dorsal striatum and spinal cord, respectively, for cell subtype identification; and (2) PFFs into the dorsal striatum for seeding αSyn pathology in the cortex. It is known that motor cortical ITNs send bilateral projections to the striatum and CSNs send collateral projections to the ipsilateral striatum (Shepherd, 2013). In this experiment, ITNs and CSNs in M2 were identified from the hemisphere contralateral and ipsilateral to striatum receiving PFFs injections, respectively. Red Retrobeads-labeled ITNs were found broadly in the layer 5 of M2 and largely overlapped with pS129-ir αSyn aggregates (**Figure 1D**). In contrast, Retrobeads-labeled CSNs were restricted in the layer 5B and nearly completely separated from the cortical subregions covered by pS129-ir αSyn aggregates (**Figure 1E**). Quantification of the proportion of Retrobeads puncta colocalized with pS129-ir aggregates (**Figure 1F, G**) showed that the percentage of pS129-ir ITNs was significantly higher than that of pS129-ir CSNs (**Figure 1H**).

Given that seeding of αSyn aggregates depends on the exposure and uptake of PFFs at axon terminals, it is likely that lack of αSyn pathology in CSNs was due to insufficient or unsuccessful uptake of PFFs by their collaterals in the dorsal striatum. To visualize corticostriatal axonal field, we unilaterally injected AAVrg-tdTomato into the striatum or the spinal cord (**Figure 2A**). As expected, ITNs axonal projections in the striatum occupied a much larger striatal subregion than that of CSNs (**Figure 2B, C**). Of note, while the PFFs injection sites stayed within the ITNs axon terminal field (**Figure 2B**), they almost completely avoided the CSNs axon terminal field in the dorsal striatum (**Figure 2C**). Such difference in striatal coverage by cortical projections may, at least partially, contribute to the different levels of αSyn aggregates in ITNs and PTNs of M2. To exclude this possibility, we injected PFFs, together with Readbeads, into the spinal cord. We found mild level of αSyn aggregates in M2 at 6 months post injections, including both cytoplasmic aggreges in the layer 5B and fibril-like aggregates in the layer 1 that were supposed to be pathology-bearing apical dendrites of CSNs (**Figure 2D**).

**Figure 2.**
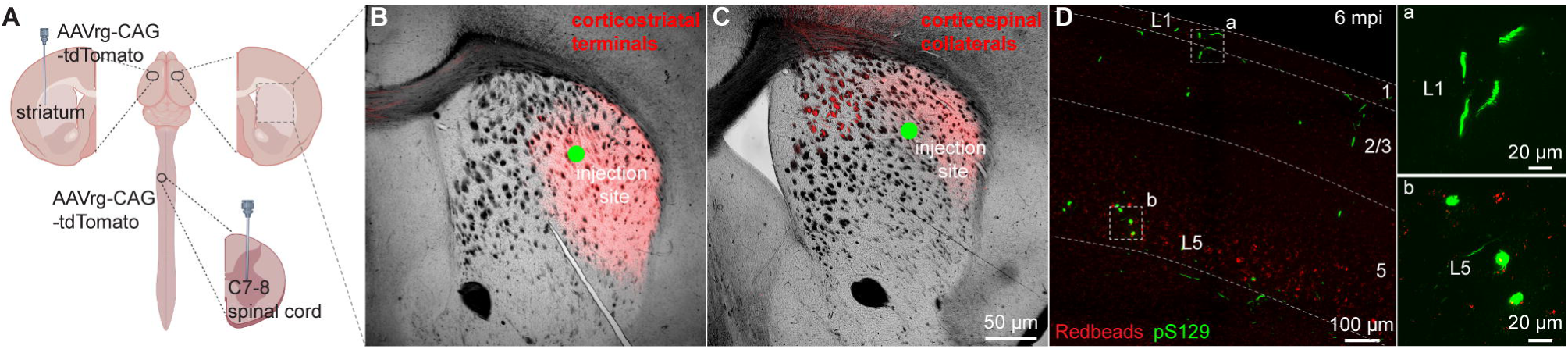
PFFs seeding into spinal cord develops moderate αSyn pathology in motor cortex. **(A)** Overall strategies to retrogradely label the ITNs and CSNs and their axonal fields in striatum. **(B and C)** Representative images showing the collateral axonal projections of ITNs (B) and CSNs (C) in striatum. The green dots indicate the approximate PFFs injection sites in the dorsolateral striatum. **(D)** Representative images showing the αSyn pathology in the motor cortex at 6 mpi time point following PFFs injection into the spinal cord. (a, b) Insets showing high magnification of αSyn pathology from the cortical regions highlighted with the squares in the layers 1 (a) and 5 (b).

Together, these results suggest that αSyn aggregates can be induced in both ITNs and CSNs of M2, but the extent of pathology seems to be greater in ITNs than that in CSNs, perhaps depending on where the seeding process starts.

### M2 neuronal hyperexcitability associated with **α**Syn aggregation

Next, we sought to understand the functional consequences of αSyn aggregation on the physiology and morphology of M2 neurons. Given their preferential accumulation of αSyn aggregates, we performed whole-cell patch-clamp recordings from retrogradely labeled M2 ITNs from the hemisphere that was contralateral to the striatum receiving PFFs injections (**Figure 3A**). We labeled all recorded neurons using biocytin through the recording pipettes for *post hoc* immunohistochemical examination of pS129-ir αSyn pathology (**Figure 3B-E**). None of the biocytin-labeled M2 neurons from PBS- or monomer-injected mice showed pS129-ir αSyn pathology (**Figure 3C**). In PFFs-injected mice, we detected somatic αSyn pathology in 14 out of 67 retrogradely labeled ITNs (i.e., pS129-positive hereafter, **Figure 3E**), but not in the other 53 cells (i.e., pS129-negative hereafter, **Figure 3D**). The small proportion of biocytin-labeled ITNs bearing αSyn aggregates is consistent with our initial observations from the histological studies (**Figure 1H**).

**Figure 3.**
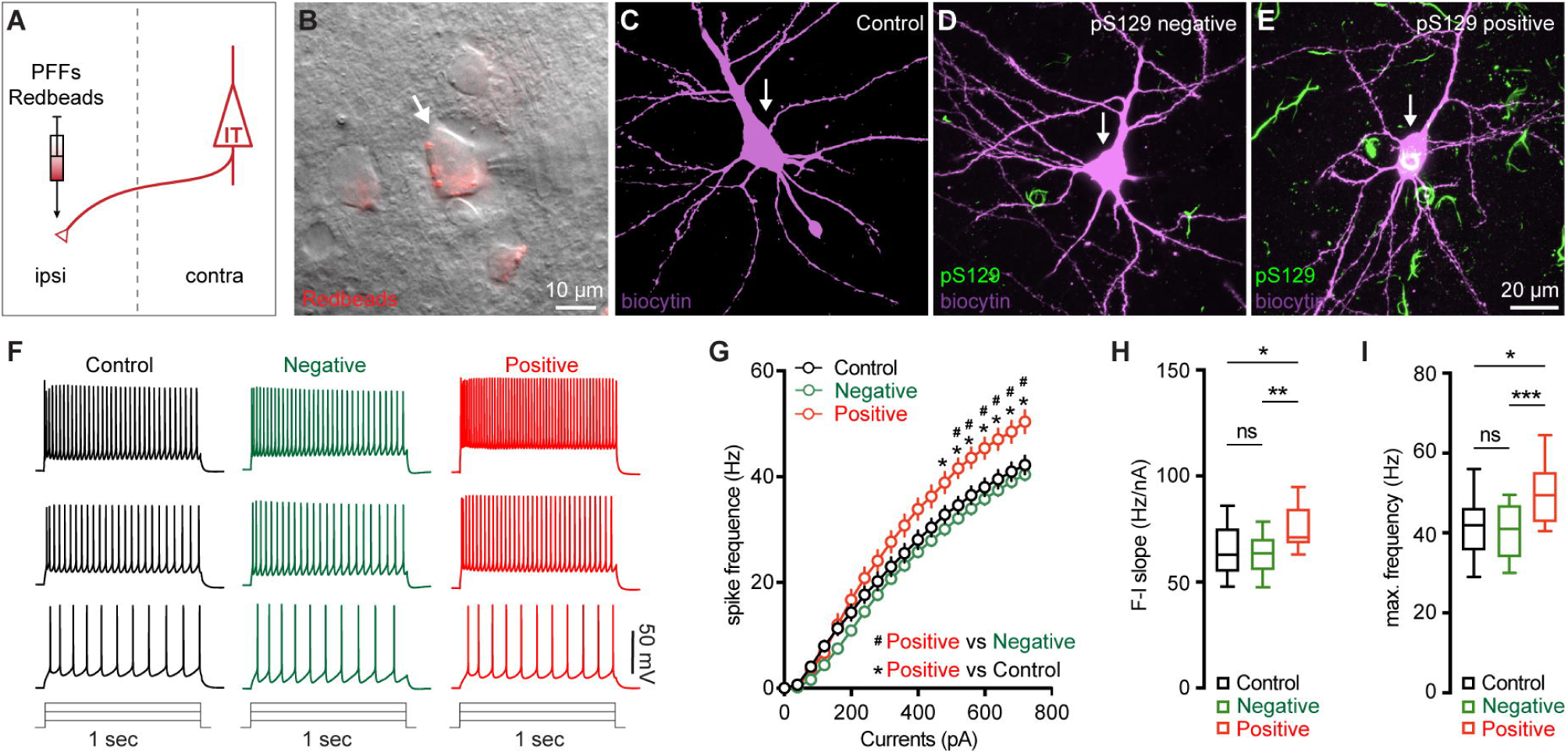
M2 neuronal hyperexcitability associated with αSyn pathology. **(A)** Diagram showing the experimental design for co-injection of αSyn PFFs with Retrobeads into the contralateral dorsal striatum. **B)** Representative images showing a Retrobeads labeled ITN in the layer 5 of M2 that was targeted for electrophysiology recording and intracellular dialysis of biocytin. **(C-E)** Representative images showing biocytin labelled M2 ITNs neurons that were from controls (C), or pS129-negative (D) and pS129-positive (green, E) from PFFs-injected mice. **(F)** Representative spike trains of ITNs evoked by somatic current injections (+160 pA, +320 pA and +480 pA for 1 sec) from controls (left), as well as pS129-negative (middle) and pS129-positive (right) cells. (**G**) Frequency-current relationship of controls as well as pS129-negative and positive cells. Controls = 34 neuron/5 mice, pS129-negative = 53 neurons/9 mice, pS129-positive = 14 neurons/9 mice. *p* < 0.0001, pS129-positive group versus controls or pS129-negative ones, mixed effects model followed by Sidak’s test. **(H-I)** Summarized results showing an increased F-I slopes (H, Controls = 34 neurons/5 mice; pS129-negative = 53 neurons/9 mice; pS129-positive = 14 neurons/9 mice; *p* = 0.0068, Kruskal-Wallis test followed by Dunn’s test) and increased maximum frequency of firing in pS129-positive group relative to controls or pS129-negative ones (I, Control = 34 neurons/5 mice; pS129-negative = 53 neurons/9 mice; pS129-positive = 14 neurons/9 mice; *p* = 0.0039, Kruskal-Wallis test followed by Dunn’s test).

After the blockade of ionotropic glutamatergic and GABAergic synaptic transmission, we injected a family of currents (1 sec) ranging from 0 to 720 pA via the patch pipettes to assess the intrinsic excitability of ITNs. We compared the intrinsic excitability of ITNs of M2 from PBS- and αSyn monomer-injected mice and found no difference in cellular excitability between two groups (**Supplementary Figure 1B-E**). Thus, we pooled cells from PBS- and monomer-injected mice as “controls” hereafter. We found that, in response to a given intensity of current injection, pS129-positive ITNs discharged more action potentials (APs) relative to pS129-negative ones in PFFs-injected mice or controls (**Figure 3F, G**). In contrast, there was no difference in the overall excitability of pS129-negative ITNs in PFFs-injected mice relative to controls (**Figure 3F, G**). We then quantified the frequency-current (F-I) curves of ITNs and found that pS129-positive ITNs exhibited a steeper F-I slop and increased maximal frequency of firing at 720 pA, relative to pS129-negative ITNs from PFFs-injected mice or controls (**Figure 3H, I**). Together the above results suggest that formation of cytoplasmic αSyn aggregates enhances the cellular excitability of ITNs in the layer 5 of M2.

### Mechanisms of M2 neuronal hyperexcitability associated with **α**Syn aggregation

To understand the ionic mechanisms underlying the hyperexcitability of M2 ITNs, we injected a family of negative current steps to assess their passive membrane properties. Membrane responses evoked by negative current injections were much greater in pS129-positive ITNs than pS129-negative ones or those from controls (**Figure 4A**). Quantitative analyses showed that pS129-positive ITNs exhibited higher input resistance and smaller cell capacitance relative to pS129-negative ITNs from PFFs-injected mice and those from controls (**Figure 4B, C**). Last, we did not detect changes in other passive membrane properties, including the resting membrane potential (Vm, **Figure 4D**), as well as the threshold, amplitude, and half-width of APs (**Figure 4E, F, G**). Together the above data show that αSyn aggregation increases the membrane input resistance, decreases cell capacitance, and enhances intrinsic excitability of cortical neurons through cell-autonomous mechanisms.

**Figure 4.**
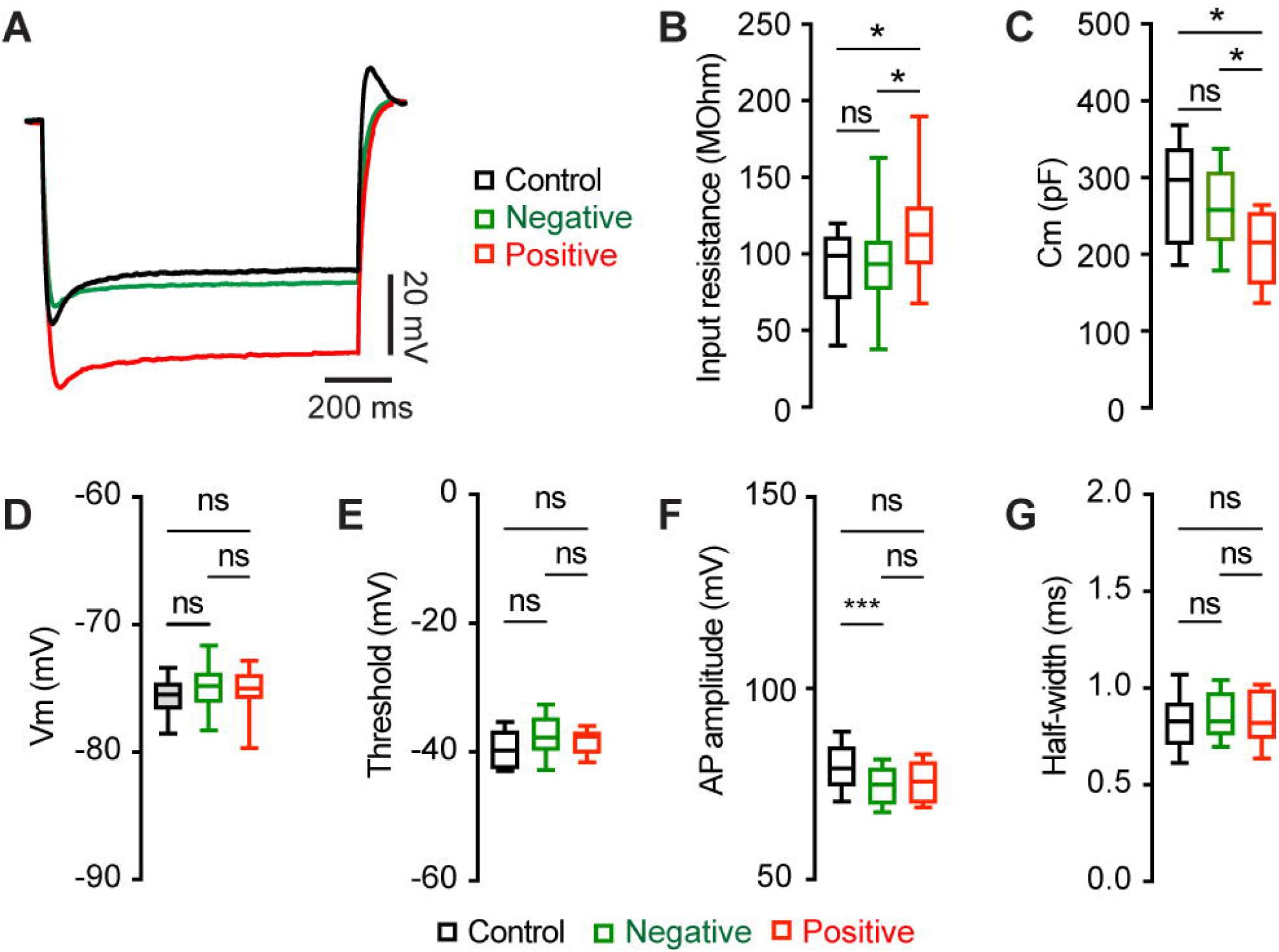
αSyn aggregation increases the input resistance of M2 ITNs. **(A)** Representative membrane responses evoked by negative current injections (-240 pA for 1 sec) of ITNs from controls and those positive and negative for pS129 aggregation. **(B and C)** Box plots showing an increased input resistance (B, Control = 93.1 [50.0, 109.0] MOhm, n = 34 neurons/5 mice; pS129-negative = 87.4 [71.2, 103.6] MOhm, n = 51 neurons/9 mice; pS129-positive = 112.7 [83.4, 136.5] MOhm, n = 14 neurons/9 mice; *p* = 0.0293) and a decreased cell capacitance (C, control = 297.2 [213.3, 337.9] pF, n = 34 neurons/5 mice; pS129-negative = 262.6 [224.5, 308.7] pF, n = 52 neurons/9 mice; pS129-positive = 215.4 [161.4, 254.3] pF, n = 14 neurons/9 mice; *p* = 0.0025) of ITNs from pS129-positive group relative to control and pS129-negative groups. **(D-G)** Boxplots showing no or subtle change in the resting membrane potential (D, control = - 75.5 [-76.6, -74.6] mV, 34 neurons/5 mice; pS129-negative = -74.8 [-76.1, -74.6] mV, n = 50 neurons/9 mice; pS129-positive = -75.0 [-76.45, -72.6] mV, n = 13 neurons/9 mice; *p* = 0.2287), threshold (E, control = -49.7 [-52.3, -46.7] mV, 33 neurons/5 mice; pS129-negative = -47.7 [-49.7, -44.7] mV, 52 neurons/9 mice; pS129-positive = -47.5 [-49.8, -46.8] mV, 14 neurons/9 mice; *p* = 0.0566), AP amplitude (F, control = 79.07 [74.63, 84.74] mV, 34 neurons/5 mice; pS129-negative = 74.86 [69.77, 79.05] mV, 52 neurons/9 mice; pS129-positive = 75.61 [70.01, 80.82], 14 neurons/9 mice; *p* = 0.0048), and AP half-width (G, control = 0.85 [0.78, 0.92] ms, n = 34 neurons/52 mice; pS129-negative = 0.83 [0.76, 0.98], n = 52 neurons/9 mice; pS129-positive = 0.82 [0.74, 0.99], n = 14 neurons/9 mice; *p* = 0.9943) of ITNs from pS129-positive group relative to controls and pS129-negative groups. Kruskal-Wallis test followed by Dunn’s test.

### Morphological changes of M2 neurons associated with **α**Syn aggregation

To further understand the impact of αSyn aggregation on cortical microcircuits, we filled Retrobeads-labeled M2 ITNs with biocytin via the patch pipettes to study potential changes in their morphology. Biocytin labelled dendritic tree and cell body were visualized by a confocal microscope, followed by a three-dimensional reconstruction for analysis. Inclusion of somatic pS129-ir αSyn aggregates were immunohistochemically examined for all M2 ITNs from PFFs-injected mice, which were then analyzed and presented separately. We found a significant reduction of dendritic arborization of pS129-positive ITNs in M2, relative to pS129-negative ITNs from PFFs-injected mice or those from controls (**Figure 5A**). Sholl analysis showed that there were significantly less dendritic branches within 200 μm from the soma of pS129-positive ITNs, relative to pS129-negative ITNs from PFFs-injected mice or controls (**Figure 5B**). A large portion of proximal dendritic branches was from the basal dendrites in the layer 5. Consistently, we found that pS129-positive ITNs had shorter length of basal dendrites compared to pS129-negative ITNs from PFFs-injected mice or controls (**Figure 5C**). Moreover, there was a significant loss of spines on the basal dendrites of pS129-positive ITNs, relative to pS129-negative ITNs from PFFs-injected mice or controls (**Figure 5D-G**). Last, we also conducted three-dimensional reconstruction of M2 ITNs and quantified the soma volume as an assessment of cell body size. We found a significant reduction in the soma size of pS129-positive ITNs of M2 relative to pS129-negative ones from PFFs-injected mice or controls (**Figure 5H**), suggesting a shrinkage of ITNs bearing αSyn aggregates, which is consistent with the described changes in cell capacitance (**Figure 4C**).

**Figure 5.**
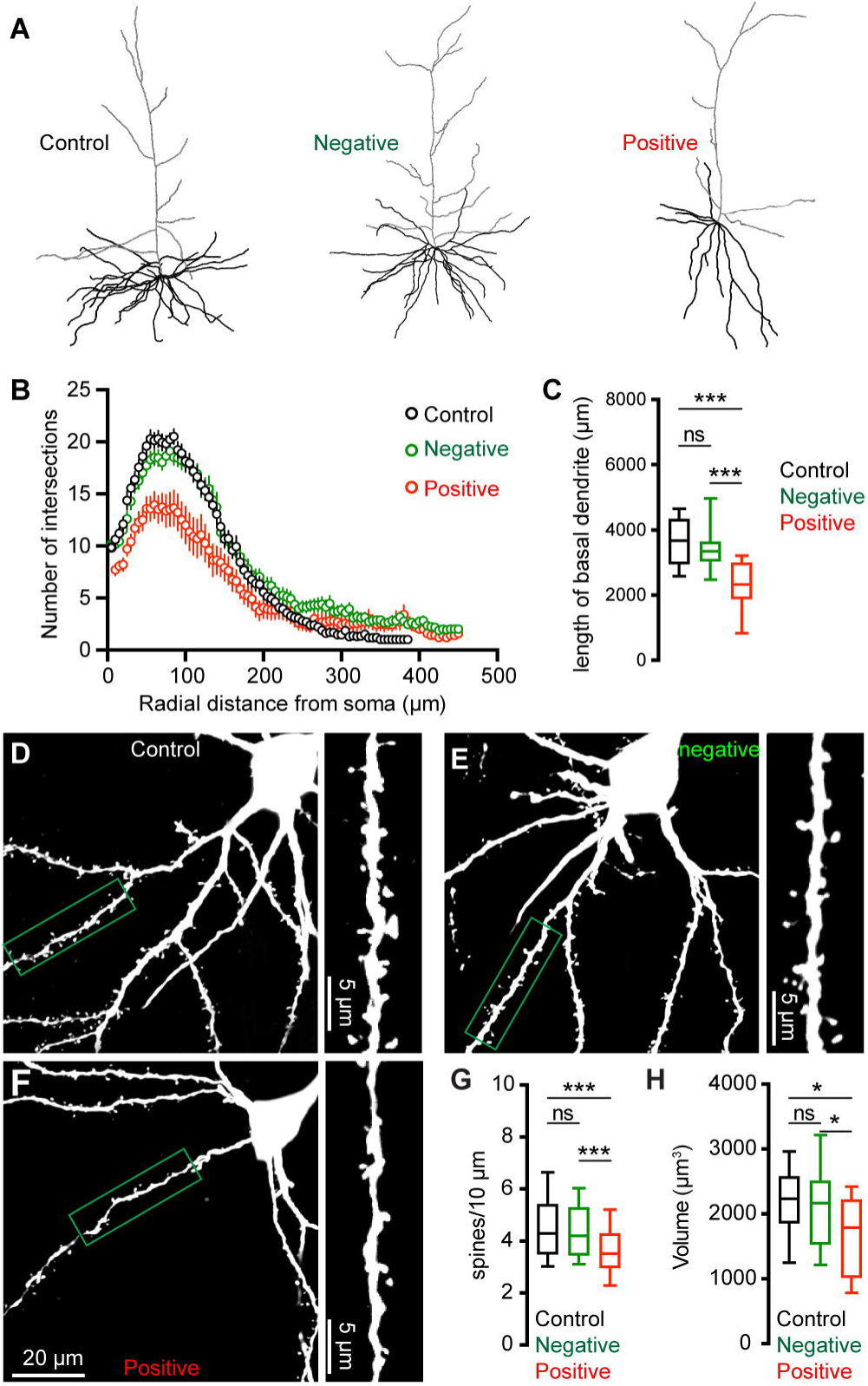
αSyn pathology induces morphological changes of ITNs. **(A)** Representative reconstructed ITNs from control, pS129-negative, and pS129-positive groups. Black and gray traces indicate basal and apical dendrites, respectively. **(B)** Sholl analysis showing a reduced dendritic branching (*p* < 0.0001, mixed effects model followed by Sidak’s test). (**C**) Box plots showing decreased total length of the basal dendrites (control = 3679 [2951, 4328] μm, n = 22 neurons/4 mice; pS129-negative = 3354 [3050, 3641] μm, 25 neurons/6 mice; and pS129-positive = 2330 [1886, 2998] μm, 14 neurons/9 mice, *p* < 0.0001) of pS129-positive ITNs relative to control and pS129-negative ones. **(D-F)** Representative images of biocytin-filled ITNs and segments of dendritic spines from control (D), pS129-negative (E), and pS129-positive (F) groups. **(G)** Box plots showing a decreased spine density in pS129-positive ITNs relative to controls and pS129-negative ones (spine density, controls = 4.290 [3.488, 5.401]/10 μm, n = 66 segments/4 mice; pS129-negative = 4.202 [3.460, 5.285]/10 μm, n = 75 segments/6 mice; pS129-positive = 3.521 [2.979, 4.297]/10 μm, n = 41 segments/9 mice; *p* = 0.0056. **(H)** Boxplots showing a decreased soma volume of pS129-positive ITNs relative to controls and pS129-negative ones. (Control = 2234 [1860, 2579] μm^3^, n = 31 neurons/5 mice; pS129-negative = 2169 [1536, 2514] μm^3^, n = 32 neurons/9 mice; pS129-positive = 1794 [1020 to 2225] μm^3^, n = 14 neurons/9 mice; *p* = 0.0331, pS129-positive versus controls; *p* = 0.0359, pS129-positive versus pS129-negative. Kruskal-Wallis test followed by Dunn’s test for C, G, H.

### αSyn aggregation does not alter motor thalamic inputs to ITNs of M2

Glutamatergic motor thalamic inputs to cortical pyramidal neurons form synapses at both basal dendrites in the layer 5 and distal apical dendrites in the layer 1 (Hooks et al., 2013; Biane et al., 2016; Villalba et al., 2019; Chen et al., 2023). The significant loss of basal dendritic spines indicates that the thalamic inputs to ITNs may be altered by αSyn aggregation. In the intrastriatal PFFs seeding model, we mixed PFFs with Retrobeads and injected the mixed solution unilaterally into the dorsal striatum for retrogradely labelling of M2 ITNs and seeding of αSyn aggregation (**Figure 6A**). The ITNs in the contralateral hemisphere, with intact nigrostriatal and mesocortical dopamine projections, were be targeted and studied. Thus the effect of αSyn aggregation was not confounded by potential impact of dopamine depletion (Chen et al., 2023).

**Figure 6.**
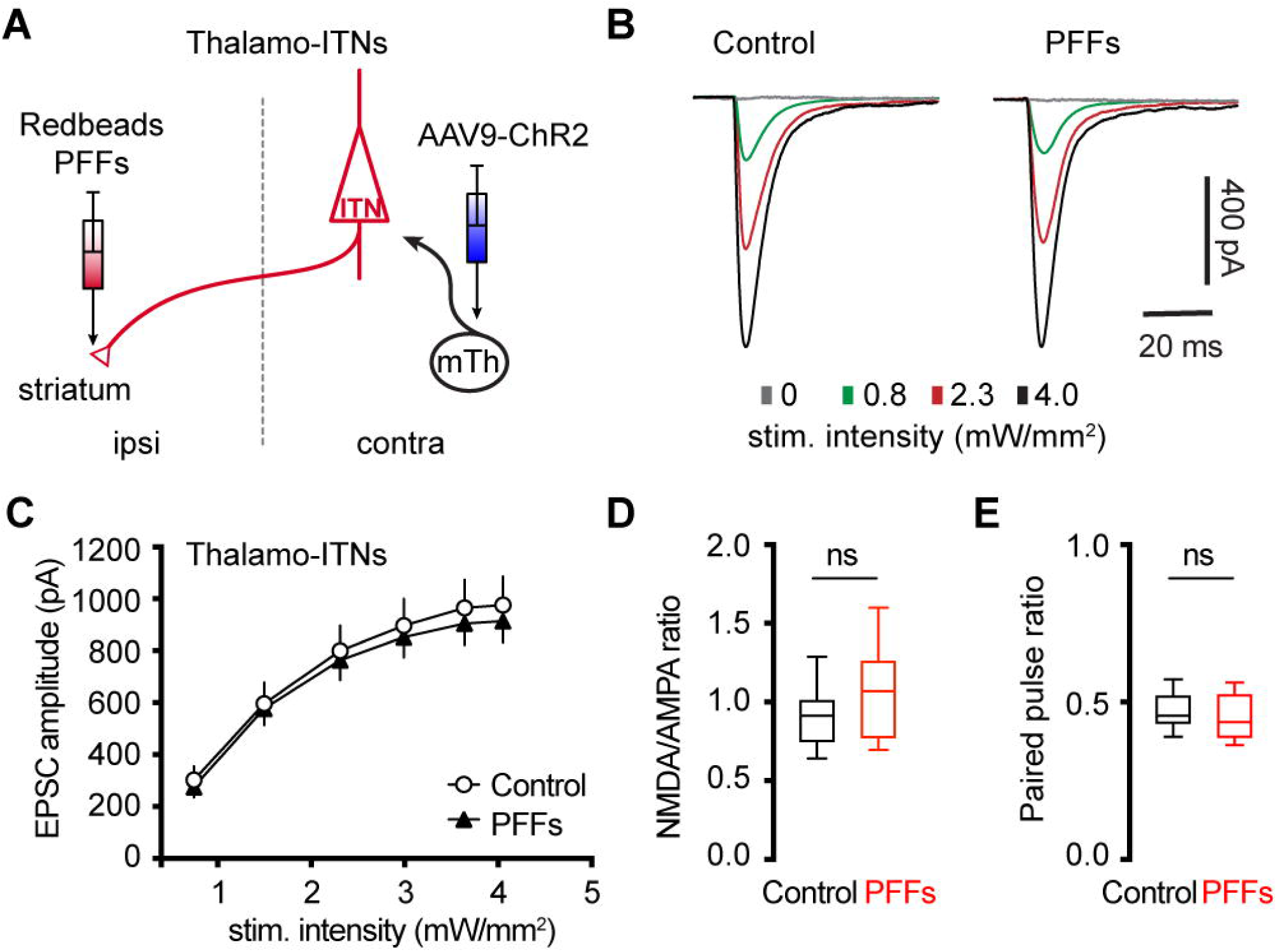
No change in the thalamocortical transmission to M2 ITNs in the PFFs seeding model. **(A)** Diagram showing the experiment design to study thalamocortical transmission of M2 ITNs. **(B and C)** Representative traces of optogenetically evoked EPSCs in ITNs across different stimulation intensities from control and PFF groups (B) and the summarized results (C, control = 26 neurons/5 mice; PFF = 18 neurons/3 mice. *p* = 0.7391, mixed effects model followed by Sidak’s tests). **(D)** Box plots showing no change in the NMDA/AMPA ratio at thalamocortical synapses to M2 ITNs between controls and PFFs-injected mice (control = 0.91 [0.75, 1.02], 26 neurons/5 mice; PFFs = 1.07 [0.77, 1.26], 18 neurons/3 mice, *p* = 0.09, MWU). **(E)** Box plots showing no change in the paired pulse ratio at thalamocortical synapses to M2 ITNs between controls and PFFs-injected mice (control = 0.46 [0.43, 0.53], 21 neurons/5 mice; PFFs = 0.44 [0.39, 0.53], 16 neurons/3 mice, *p* = 0.44, MWU).

To interrogate the synaptic strength of thalamic inputs to the ITNs of M2, we injected AAV9-ChR2(H134R)-eYFP into the motor thalamus that was contralateral to the hemisphere receiving PFFs/Retrobdeas injections (**Figure 6A**). Upon stimulation of ChR2-expressing thalamic axon terminals in M2 by delivering 478 nm light, robust excitatory postsynaptic currents (EPSCs) were recorded using whole-cell voltage-clamp recording at -80 mV, which could be completely abolished by AMPA receptor antagonist DNQX (20 μM). A range of intensities of blue light were delivered to assess the connection strength of thalamo-ITNs synapses in both controls and PFFs-injected mice. We found that there was no change in the amplitude of thalamic EPSCs in ITNs from PFFs-injected mice relative to those from controls across a range of light intensities (**Figure 6B, C**). Pathological αSyn aggregation may interact with postsynaptic NMDA receptor subunits, as reported in the striatum (Tozzi et al., 2016). To test this possibility in the cortical circuits, we quantified the ratio of NMDA- and AMPA-receptor mediated EPSCs (NMDA/AMPA ratio) at thalamo-ITNs, and found no difference in the NMDA/AMPA ratio between controls and PFFs-injected mice (**Figure 6D**). Furthermore, we did not detect change in paired pulse ratios (PPR) at the thalamo-ITNs between PFFs-injected mice and controls (**Figure 6E**), indicating a lack of alterations of the initial presynaptic release probability following αSyn pathology formation. These data were supported by an absence of prominent αSyn pathology accumulation in the ventromedial region of the motor thalamus (**Supplementary Figure 2**). The above results are consistent with recent studies showing low levels of *SNCA* gene expression in the motor thalamus and αSyn protein expression in vGluT2-expressing thalamic neurons and their axon terminals in the cerebral cortex (Chen et al., 2022; Geertsma et al., 2024).

### Partial dopamine depletion does not alter the excitability and synaptic excitation of corticospinal neurons in M2

At 3 months post injections, moderate levels of pS129 αSyn pathology accumulated in the ventral tier of the substantia nigra compacta of PFFs-injected mice, but no obvious cell loss was detected (**Supplementary Figure 3A, B**). Consistently, the density of TH immunoreactive axon terminals in the ipsilateral striatum decreased ∼30% in PFFs injected mice (**Supplementary Figure 3E**). We recently reported an impaired intrinsic excitability and reduced thalamic excitation to motor cortical PT neurons in parkinsonian mice with nearly complete dopamine depletion (Chen et al., 2021; Chen et al., 2023). Therefore, although corticospinal neurons did not develop detectable αSyn aggregation in the intrastriatal PFF model (**Figure 1 and Figure 7C**), their intrinsic and synaptic properties may be affected by the partial striatal dopamine loss and the associated basal ganglia dysfunction (Escande et al., 2016; Willard et al., 2019).

**Figure 7.**
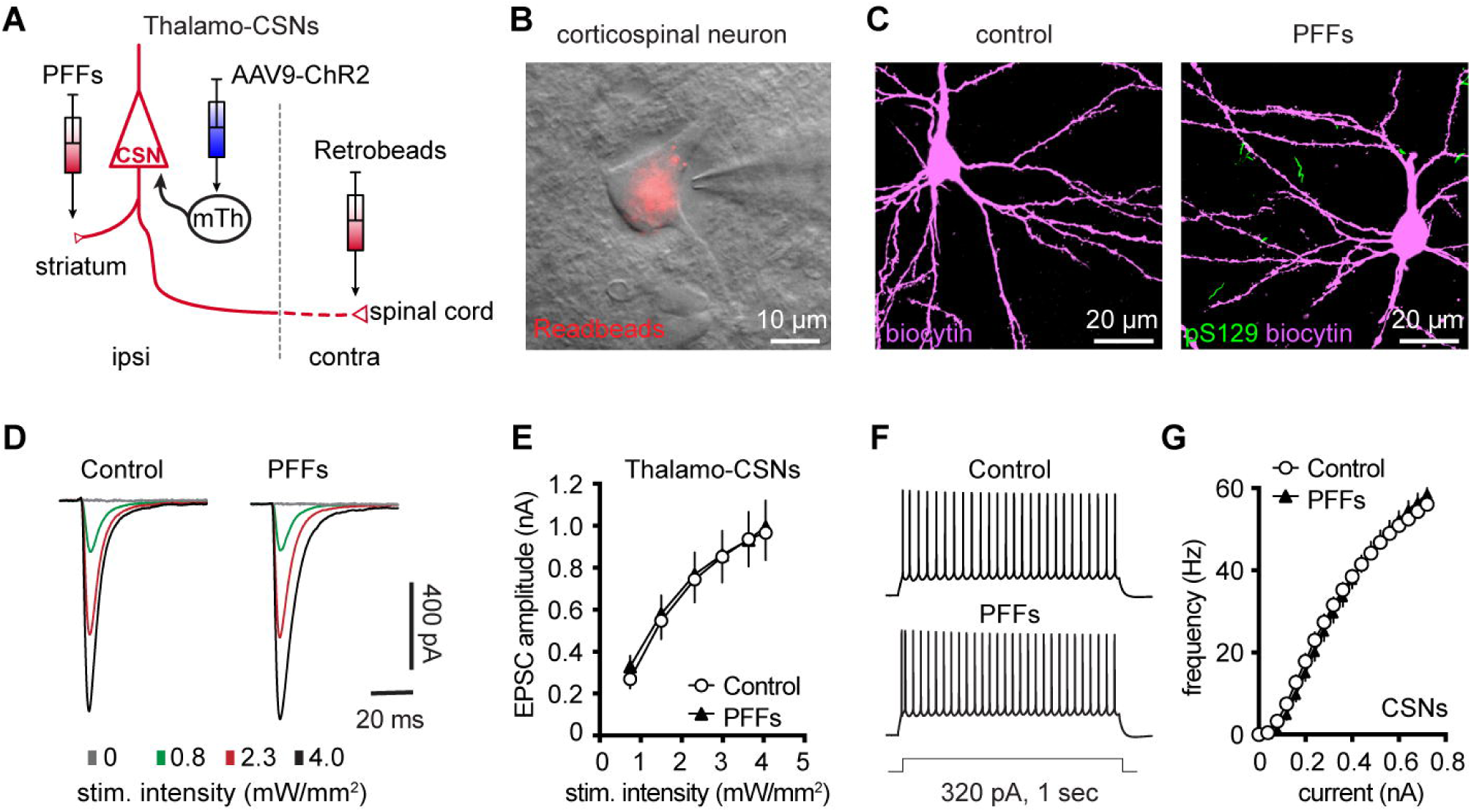
Intrinsic and synaptic properties of M2 CSNs were not affected in the inrtrastriatal PFFs-seeding model of synucleinopathy. **A)** Experimental design to study intrinsic and synaptic properties of corticospinal neurons in the intrastriatal PFFs-seeding model using optogenetic approaches. **B**) Representative image of a Retrobeads labeled corticospinal neuron that was targeted for physiological recording and dialysis of biocytin. **C**) Representative confocal images showing biocytin-filled corticospinal neurons from controls and PFFs-injected mice. Both neurons were negative of pS129 immunoreactivity in post hoc IHC staining. **D**) Representative traces of optogenetically evoked EPSCs in CSNs across different stimulation intensities from controls and PFFs-injected mice. (**E**) Summarized results showing no change in the amplitude of thalamocortical EPSCs in CSNs between controls and PFFs-injected mice (controls = 24 neurons/5 mice; PFF = 24 neurons/4 mice. *P* = 0.8605, mixed effects model followed by Sidak’s tests). **F**) Representative AP spike trains of CSNs from controls and PFFs-injected mice in response to somatic current injections (320 pA for 1 sec). **G**) Summarized graph showing no changes in the excitability of CSNs from controls and PFFs-injected mice (*p* = 0.8110, Kruskal-Wallis test. control = 28 neurons/5 mice; PFF = 20 neurons/3 mice).

To test this hypothesis, we injected (1) αSyn PFFs into the dorsal striatum to induce the formation of αSyn aggregation in the SNc dopaminergic neurons and partial striatal dopamine depletion (**Supplementary Figure 3B-E**); (2) AAV9-ChR2(H134R)-eYFP into the motor thalamus of the same hemisphere as the striatum receiving PFFs injections; and (3) Retrobeads to the spinal cord to retrogradely label corticospinal neurons for electrophysiology recordings (**Figure 7A**). Upon optogenetic stimulation of ChR2-expressing thalamic axon terminals in M2, we detected robust thalamic EPSCs in corticospinal neurons, as previously reported (Hooks et al., 2013; Biane et al., 2016; Chen et al., 2023). At 3 mpi, we did not detect difference in the amplitudes of thalamic EPSCs in corticospinal neurons between controls and PFFs injected mice (**Figure 7D, E**), suggesting that there was no change in the connection strength of thalamocortical inputs to the corticospinal neurons. Thus, we concluded that a partial loss of striatal dopamine does not affect the connection strength of thalamic inputs to corticospinal neurons.

Furthermore, we assessed the intrinsic excitability of M2 corticospinal neurons by performing whole-cell current-clamp recordings from PFFs-injected mice and controls at 3 mpi. We found that corticospinal neurons discharge similar number of APs in response to a range of current injections between groups (**Figure 7F, G**). These results suggest that the intrinsic excitability of M2 corticospinal neurons was not altered by a partial dopamine depletion in the intrastriatal PFFs model.

## Discussion

The present study demonstrated that (1) formation of intracellular αSyn aggregates increases the cellular excitability of M2 pyramidal neurons, (2) the increased intrinsic excitability likely occurs through cell autonomous mechanisms; and (3) a partial degeneration of nigrostriatal pathway is not sufficient to trigger adaptative changes of the intrinsic and synaptic properties of M2 corticospinal neurons. Taken together, the present study provides novel insights into cortical pathophysiology in parkinsonism.

Post-mortem analyses of brains of PD patients and animal models have identified a group of brain regions that show particular vulnerability to accumulation of αSyn aggregations, including the SNc, locus coeruleus, dorsal vagal nucleus, among many others (Braak et al., 2003). Compelling evidence suggests that αSyn pathology propagates through brain networks, driving neuronal death and loss of synaptic connections within and between the vulnerable brain regions and cell subtypes (Kordower et al., 2008; Li et al., 2008; Osterberg et al., 2015; Surmeier et al., 2017). However, not all cells bearing αSyn pathology degenerate in brain, indicating not only a poor correlation between Lewy pathology and cell death but also critical roles of cell intrinsic mechanisms in gating neurodegeneration (Surmeier et al., 2017). In addition, detrimental consequence of αSyn aggregation to neuronal and synaptic function could be the major driver of phenotype manifestation in PD and other synucleinopathies (Kulkarni et al., 2022). Considering the preferential presynaptic location of αSyn, a large body of studies have focused on studying the effects of its abnormal aggregation to synaptic structure and function (Volpicelli-Daley et al., 2011; Nakata et al., 2012; Vargas et al., 2014). These reports highlight disrupted synaptic transmission at both dopaminergic and glutamatergic systems that may underlie either motor and nonmotor deficits associated with αSyn aggregation (Janezic et al., 2013; Tozzi et al., 2016; Phan et al., 2017; Wu et al., 2019; Tozzi et al., 2021).

On the other hand, emerging evidence suggests that abnormal aggregation of αSyn triggers adaptive changes in intrinsic excitability of dopaminergic neurons in the SNc and cholinergic neurons in the vagal nucleus (Subramaniam et al., 2014; Lasser-Katz et al., 2016; Chiu et al., 2021; Ledonne et al., 2023). Our studies further expand these series of observations to the motor cortical glutamatergic circuits, which exhibit Lewy pathology at late stages in PD. A key finding of our study is that αSyn aggregates-bearing cortical pyramidal neurons showed increased excitability relative to those without cytoplasmic αSyn aggregates from the PFFs-injected mice or controls (**Figure 3**). Since the intrinsic excitability was assessed in the presence of ionotropic glutamatergic and GABAergic receptors, the hyperexcitability of pathology-bearing cortical neurons was likely to be mediated by cell-autonomous mechanisms. These data provide mechanistic understanding of cortical hyperexcitability observed *in vivo,* which was initially explained mainly by a disrupted balance between synaptic excitation/inhibition (Blumenstock et al., 2021; Ramalingam et al., 2023). Mechanistically, we showed that the hyperexcitability of M2 pyramidal neurons was associated with an increased input resistance, decreased cell capacitance, and shrinkage of cell bodies (**Figures 4 and 5**). These data suggest that the hyperexcitability of αSyn aggregates-bearing cells might be due to abnormal membrane expression and re-distribution of ion channels (e.g., voltage-gated Ca^2+^ and K^+^ channels) as reported in the SNc dopaminergic neurons and vagal cholinergic neurons (Subramaniam et al., 2014; Lasser-Katz et al., 2016; Chiu et al., 2021; Chiu et al., 2024).

Due to the low proportion of cortical pyramidal neurons bearing intracellular αSyn aggregates in the PFFs-seeding model (**Figure 1**) and the absence of online markers to visualize these cells, it is not technically feasible to investigate ionic mechanisms underlying the hyperexcitability of cortical pyramidal neurons using pharmacological or genetic approaches. In addition to cell-autonomous adaptations, we cannot exclude potential contribution of non-cell autonomous mechanisms to neuronal and network hyperexcitability *in vivo*, such as the involvement of glial activation and neuroinflammatory responses (Umpierre and Wu, 2021; Targa Dias Anastacio et al., 2022).

Though M2 ITNs showed hyperexcitability, it does not necessarily mean their output to the striatum and other subcortical regions is enhanced. Contrarily, evidence in the literature suggests a significant reduction of cortical glutamatergic outputs in model of synucleinopathies that may occur at a relatively earlier time point (Wu et al., 2010; Guatteo et al., 2017; Chen et al., 2022).

Cortical ITNs exhibited shrinkage of cell bodies and loss of dendritic spines following the formation of intracellular αSyn aggregation (**Figure 5**), which were similar to the morphological changes in the SNc dopamine neurons and vagal cholinergic neurons (Chiu et al., 2021; Ledonne et al., 2023). Surprisingly, loss of spines of basal dendrites was not associated with alterations of thalamocortical connection strength of ITNs and presynaptic release probability in the M2 (**Figure 6**). Given that detrimental effect of αSyn aggregation to presynaptic transmission (Volpicelli-Daley et al., 2011; Vargas et al., 2017), this observation is consistent with relatively low expression levels of endogenous αSyn in most thalamic subregions (Taguchi et al., 2015; Chen et al., 2022), and is supported by the absence of pathological αSyn aggregates in the ventromedial thalamus. However, a limitation of the present study was the lack of distinguishing ITNs based on the absence and presence of αSyn aggregates, which might result in underestimated postsynaptic effects of αSyn aggregation to thalamocortical synaptic connection.

Motor cortical PTNs are preferentially and severely affected by midbrain dopaminergic neurodegeneration in parkinsonism (Goldberg et al., 2002; Pasquereau and Turner, 2011). Consistently, we recently reported that both the intrinsic excitability and thalamocortical transmission were selective downregulated in PTNs, but not ITNs, using mice with 6-hydroxydopamine lesion (Chen et al., 2021; Chen et al., 2023). Further analysis indicated that postsynaptic NMDA receptors of pyramidal tract neurons may play a critical role in mediating cortical circuit adaptations in parkinsonism. In the present study, we further demonstrated that a partial loss of striatal dopamine had no effect to the intrinsic excitability and thalamocortical transmission of cortical corticospinal neurons (**Figure 7**). These data suggest that striatal dopamine loss has to be greater than a certain threshold to trigger NMDA receptors-mediated cortical circuit adaptations. A gradual development of synchronized bursting pattern of activity throughout the basal ganglia-thalamocortical network might be a key network determinant in mediating an effective stimulation of postsynaptic NMDA receptors in cortical pyramidal neurons (Bruno and Sakmann, 2006). Of particular interest, the emergence of the synchronized bursting pattern of network activity involves neural plasticity processes associated with chronic and robust dopamine depletion in the SNc (Vila et al., 2000; Mallet et al., 2008; Willard et al., 2019). Thus, motor cortex may exhibit secondary circuit adaptations at late stages of nigrostriatal dopamine neurodegeneration, as a consequence of pathological basal ganglia outputs to the motor thalamus.

αSyn pathology may reach the cerebral cortex at the Braak stage 4 and beyond (Braak et al., 2003; Fu et al., 2022). It is important to study whether αSyn pathology and nigral dopaminergic degeneration interact to disrupt the integrity and function and cortical circuits in PD. The present study presented preliminary data, as an initial attempt to address this question. Detailed analysis of cortical circuits in the presence of both severe dopamine depletion and Lewy-like pathology remains needed to advance our understanding of cortical circuit operation under parkinsonian state.

## Acknowledgement

Experimental work was conducted at Van Andel Research Institute. Data analysis was partially performed at Van Andel Research Institute and completed at Georgetown University Medical Center. The authors thank Drs. Vijay Singh and Marissa Menard in Dr. Volpicelli-Delay lab at the University of Alabama at Birmingham (UAB) for generation and validation of mouse α-synuclein preformed fibrils. We thank Dr. Laura Volpicelli-Daley for sharing mouse PFFs and thoughtful comments on this manuscript. The authors thank Dr. Thomas Wichmann at Emory University for the constructive comments on this work. This work was partially supported by research grants from National Institutes of Health (R01NS121371, H.Y.C), Department of Defense Congressionally Directed Medical Research Programs (W81XWH-21-1-0943, H.Y.C.), and the MIND program from the Van Andel Institute (L.C.). This work was also funded in whole or in part by Aligning Science Across Parkinson’s (ASAP-020572) through the Michael J. Fox Foundation for Parkinson’s Research (MJFF). For the purpose of open access, the authors have applied a CC BY public copyright license to all Author Accepted Manuscripts arising from this submission.

## Availability of data and materials

The datasets used and/or analyzed during the current study have been deposited and are publicly available from https://doi.org/10.5281/zenodo.13323976.

## Authors’ contributions

LC and HDC conducted stereotaxic surgeries and confocal imaging; LC performed electrophysiological recordings and data analysis, as well as wrote initial draft. HYC contributed to conceptualization of the project, obtaining grant supports and the administration, as well as analyzing and interpreting data. All authors contributed to writing, editing and revising the manuscript, and approved the final manuscript.

