## Supplementary figures and images for "Motor Cortical Neuronal Hyperexcitability Associated with α-Synuclein Aggregation"

### Supplement figure 1

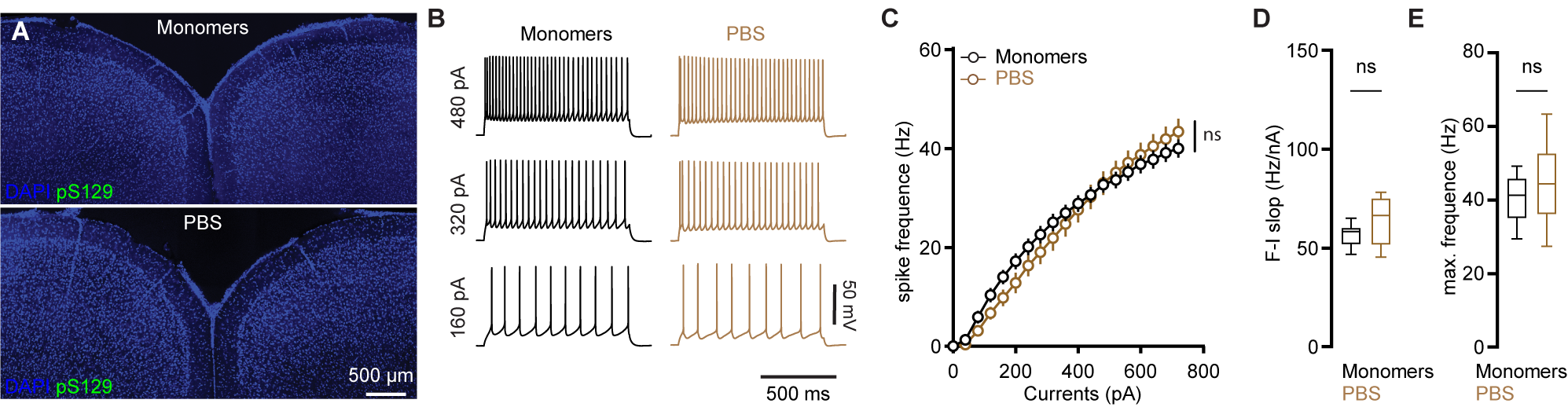

### Supplement figure 2

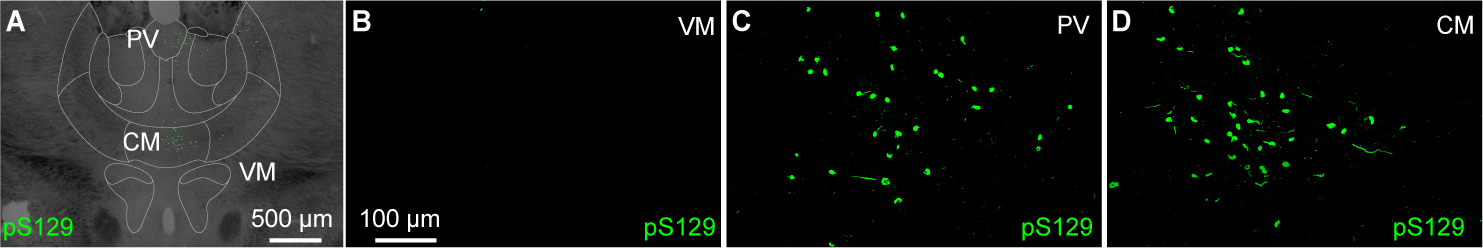

### Supplement figure 3

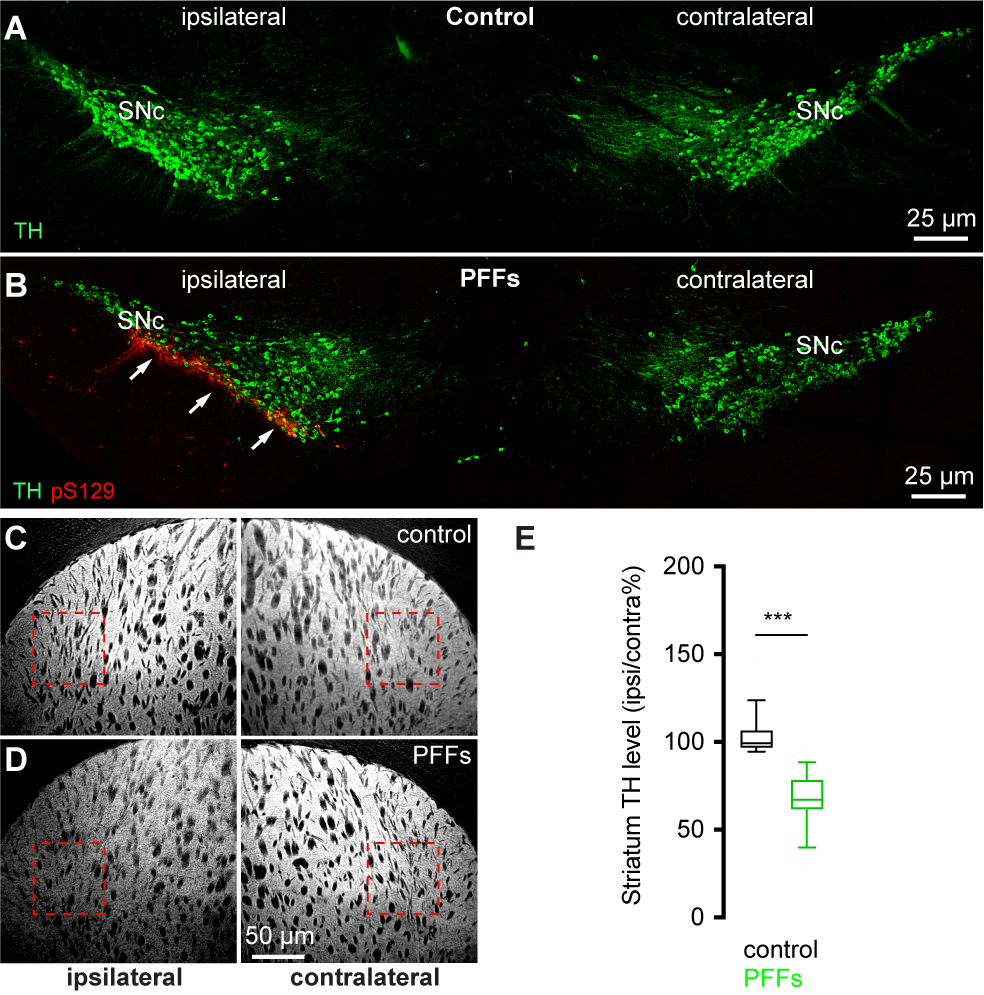
